# Decellularized Articular Cartilage Microparticles for Expansion of Mesenchymal Stem Cells and Zonal Regeneration of Articular Cartilage

**DOI:** 10.1101/2021.04.16.440121

**Authors:** Azadeh Sepahvandi, Safaa Ibrahim Kader, Mehri Monavarian, Victor Anthony Madormo, Esmaiel Jabbari

**Author notes:** The first and second authors contributed equally to this manuscript. **Corresponding author:** Esmaiel Jabbari, Ph.D., Full Professor of Chemical and Biomedical Engineering, Department of Chemical Engineering, Swearingen Engineering Center, Rm 2C11, University of South Carolina, Columbia, SC 29208.

## Abstract

**Introduction:** The objective was to create multilayer cellular constructs using fetal or adult, decellularized articular cartilage in particulate form as microcarriers for expansion and fusion of mesenchymal stem cells (MSCs) to regenerate the stratified structure of articular cartilage.

**Methods:** Porous microparticles (CMPs) generated from decellularized fetal or adult bovine articular cartilage were used as microcarriers for expansion of human MSCs. The CMP expanded MSCs (CMP-MSCs) were used to generate injectable hydrogels or preformed multilayer constructs for articular cartilage regeneration. In the injectable approach, CMP-MSCs were suspended in alginate gel, crosslinked with calcium chloride, and incubated in chondrogenic medium to generate an injectable regenerative construct. In the preformed approach, fetal or adult CMP-MSCs were suspended in a culture medium, allowed to settle sequentially by the force of gravity, and fused by incubation in chondrogenic medium to generate multilayer cell sheets. The constructs were characterized with respect to compressive modulus, cellularity, and expression of chondrogenic markers.

**Results:** Human MSCs expanded on fetal or adult CMPs in basal medium maintained the expression of mesenchymal markers. The injectable CMP-MSCs hydrogels had significantly higher expression of chondrogenic markers and compressive modulus after four weeks incubation in chondrogenic medium compared to MSCs directly encapsulated in alginate gel; preformed CMP-MSCs cell sheets had significantly higher compressive modulus and expression of chondrogenic markers compared to MSCs in the pellet culture.

**Conclusion:** The preformed cell sheet approach is potentially useful for creating multilayer constructs by sequential gravitational settling of CMP-MSCs to mimic the stratified structure of articular cartilage.

**Insight, Innovation, Integration:** This work described a novel approach to recreate the zonal structure of articular cartilage. Human MSCs were expanded on porous microcarrier beads generated from decellularized fetal or adult bovine articular cartilage. The cell-seeded microbeads were fused by gravitational settling to form mono- or bi-layer cell sheets. The cell sheets were cultured in chondrogenic medium to regenerate the articular cartilage tissue. The *in vitro* regenerated tissue had higher compressive modulus and expression of chondrogenic markers compared to the MSC pellet culture.

## 1. Introduction

Cartilage degeneration is prevalent in older adults with 65% of those over 60 years old experiencing joint pain and long-term disability [1–3]. Due to its complex physical and biochemical properties, there has been limited clinical success in treatment of articular cartilage injuries [4]. Autologous Chondrocyte Implantation (ACI) is currently used in the clinic to treat cartilage injuries. Although ACI shows better long-term outcomes than microfracture technique or other traditional methods [5, 6], its success is limited by poor cell retention, non-homogenous cell distribution, dedifferentiation of implanted cells, periosteal hypertrophy, and donor site morbidity [7, 8]. Thus, alternative methods have been explored for the repair of cartilage defects.

A promising alternative to chondrocyte harvesting is the use of adult human mesenchymal stem cell, hereafter referred to as MSCs, derived from the bone marrow or synovium [9, 10]. MSCs delivered in a supportive matrix have been shown to promote the expression of chondrogenic markers and produce a cartilage-like matrix *in vivo*. However, the approach of MSC encapsulation in a uniform matrix, without gradients, often leads to fibrocartilage formation and tissue degeneration [11, 12]. The formation of inferior fibrocartilage tissue is rooted in the inability of cellular constructs to recapitulate the stratified structure of articular cartilage [13]. There is clearly a need for novel engineering approaches to recreate the zonal structure of full-thickness articular cartilage defects without fibrocartilage tissue formation.

The stratified structure of articular cartilage is composed of the superficial, middle, deep and calcified zones with each zone having a defined protein expression, cellularity, extracellular matrix (ECM) composition and structure for lubrication, compressive strength, and load transfer to the subchondral bone [14]. It is well established that fetal articular cartilage, with a stratified structure from week 12 of gestation, has a higher regenerative capacity compared to the adult, mainly due to difference in cellularity and ECM composition [15]. Engineering approaches that mimic cellularity, ECM composition and structure of fetal articular cartilage could enhance regeneration of full-thickness articular cartilage defects.

We previously demonstrated for the first time that high cellularity, low matrix stiffness and combination of transforming growth factor-β1 (TGF-β1) and bone morphogenetic protein-7 (BMP-7) led to chondrogenic differentiation of MSCs to the superficial zone phenotype of articular cartilage; medium cellularity and stiffness, and combination of TGF-β1 and IGF-1 led to the middle zone phenotype; and low cellularity, high matrix stiffness and combination of TGF-β1 and hydroxyapatite (HA) led to the calcified zone phenotype [16, 17]. Further, we recently demonstrated that MSCs encapsulated in digested, decellularized articular cartilage could sequentially be differentiated to the superficial zone phenotype, followed by the middle and calcified zones by sequential supplementation of the chondrogenic medium with BMP-7, insulin growth factor-1 (IGF-1) and Indian hedgehog (IHH) [18]. In addition, we showed that digested and decellularized fetal articular cartilage induced differentiation of MSCs to the superficial zone phenotype of chondrocytes whereas the adult cartilage induced differentiation to the calcified phenotype.

Biodegradable microcarriers, due to their large surface area, have been used as 3D matrices for expansion of stem cells [19, 20]. Commonly used microcarriers, aside from triggering undesirable phenotypic changes, require additional processing steps to separate the expanded cells from the carrier for clinical applications. We hypothesized that MSCs attached to developmentally inspired, articular cartilage microparticles (CMPs) can mimic the process of fetal development when combined with zone-specific growth factors to regenerate the stratified structure of articular cartilage [21]. Multilayer constructs with gradients in cell density, matrix composition, and morphogens can be assembled with from MSCs attached to CMPs, hereafter referred to as CMP-MSCs, for articular cartilage regeneration.

The following approach, illustrated schematically in Figure 1, was used to test the hypothesis. Fetal or adult bovine articular cartilage was decellularized, milled in liquid nitrogen, and the freeze-dried fragments were grinded and sorted to generate fetal (fCMPs) or adult (aCMPs) CMP fractions of different average sizes. Next, the fetal or adult MSCs were used as microcarriers for expansion of MSCs to generate fetal or adult CMP-MSCs, respectively. Then, an injectable and a prefabricated approach were used to generate articular cartilage tissues. In the injectable approach, fetal or adult CMP-MSCs were encapsulated in alginate hydrogel, hereafter referred to as CMP-MSCs/alg, and cultured in chondrogenic medium to form cartilage tissues. In the prefabricated approach, fetal or adult CMP-MSCs were suspended in chondrogenic medium, allowed to settle gravitationally to the bottom surface of the culture plate, and incubated in chondrogenic medium to form a continuous cell monolayer sheet (CMP-MSCs/ml). Fetal and adult bilayer cell sheets (CMP-MSCs/bl) were generated by gravitational settling and incubation of adult CMP-MSCs to form an adult cell sheet followed by gravitational settling and incubation of fetal CMP-MSCs on top of the adult cell sheet. The CMP-MSCs/alg, CMP-MSCs/ml and CMP-MSCs/bl were assessed with respect to compressive modulus and expression of chondrogenic markers of the superficial (Sox-9 and SZP), middle (Col II and AGC), and calcified (Col X and ALP) zones of articular cartilage.

**Figure 1.**
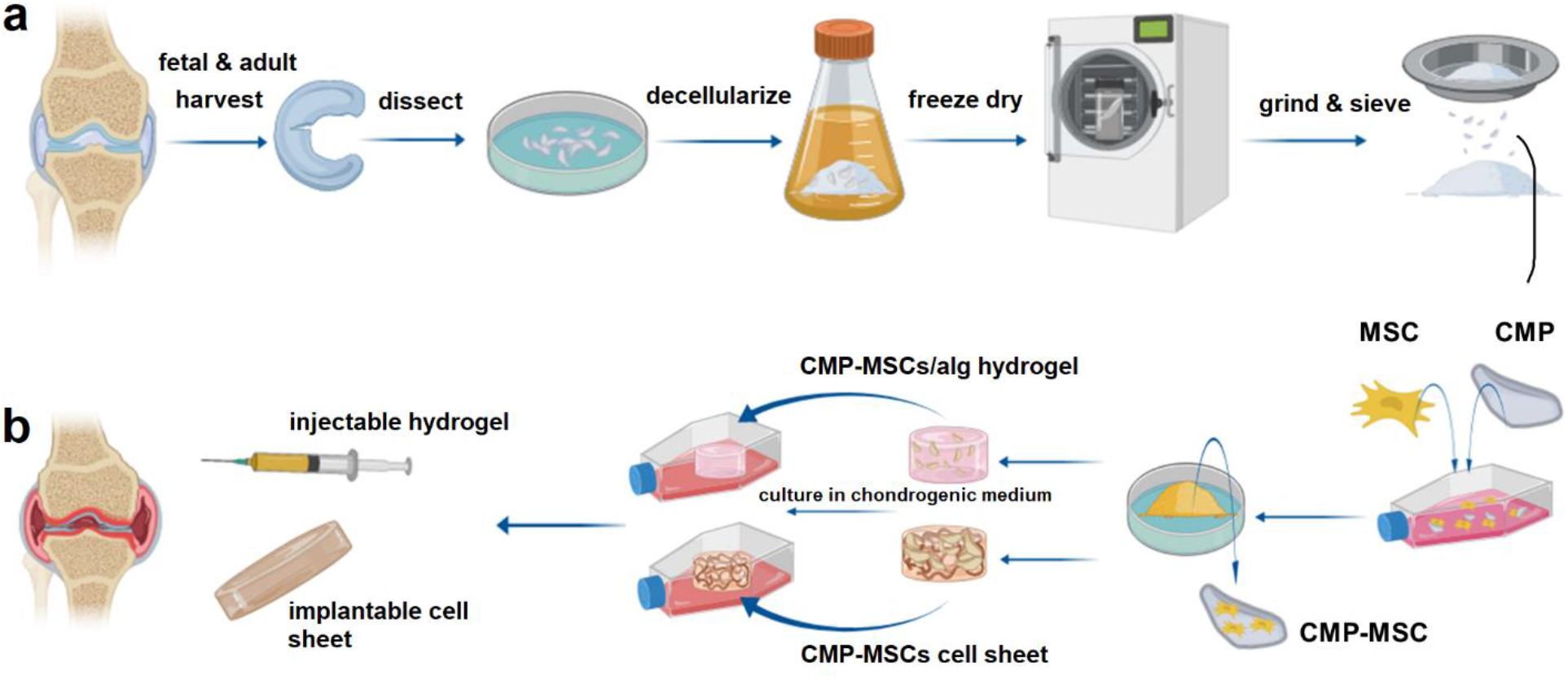
Schematic of the process for production of injectable or preformed CMP-MSCs: (a) fetal or adult articular cartilage was harvested from bovine cadaver, minced, decellularized, freeze-dried, grinded in liquid nitrogen, sieved to produce fetal or adult cartilage microparticles (CMPs); (b) MSCs were seeded on the CMPs and expanded in a tissue culture bioreactor to form CMP-MSCs. In one approach, the CMP-MSCs were suspended in alginate gel and crosslinked with calcium chloride to generate an injectable hydrogel for articular cartilage regeneration. In another approach, the CMP-MSCs suspended in medium were sequentially settled by the force of gravity on the culture plate, and incubated to generate preformed, multilayer cell sheets.

## 2. Materials and Methods

### 2.1. Materials

Paraformaldehyde, formalin, paraffin, ethylenediaminetetraacetic acid (EDTA), penicillin G, streptomycin and tris(hydroxymethyl)aminomethane (Tris) were received from Sigma-Aldrich (St. Louis, MO). The sodium salt of pyruvic acid, L-proline, DNase I and RNase A were received from Amresco (Dublin, Ireland). Sodium alginate was received from Ward’s Science (Henrietta, NY). Iodoacetic acid and calcium chloride were received from ThermoFisher Scientific (Rockford, IL). Dichloromethane (DCM, Acros Organics, Pittsburg, PA) was dried by distillation over calcium hydride. Diethyl ether, dimethylformamide (DMF) and hexane were received from VWR (Bristol, CT) and used as received. Polyester filter sieves with 80, 90, 180, 190, 240 and 250 μm pore size were obtained from Rosin Tech (Los Angeles, CA). Nylon cell strainers with 40 and 70 μm sizes were received from Corning (Canton, NY). Polystyrene microparticles with 50 μm diameter without surface modification were received from Advance Scientific (Moffat Beach, QLD, Australia).

Full-thickness adult and fetal articular cartilage harvested from bovine femoral condyles were received from Animal Technologies (Tyler, TX). Adult human mesenchymal stem cells (MSCs) harvested from healthy human bone marrow with high expression of CD105, CD166, CD29, and CD44 and low expression of TGF-β1 and CD14, CD34 and CD45 markers were received from Lonza (Allendale, NJ). Dulbecco’s modified eagle medium (DMEM), Dulbecco’s phosphate-buffer saline (PBS), fetal bovine serum (FBS), trypsin-EDTA, Quant-it PicoGreen dsDNA reagent kit, and live/dead staining kit consisting of acetomethoxy derivative of calcein (cAM) and ethidium homodimer (EthD) were received from Life Technologies (Grand Island, NY). QuantiChrom alkaline phosphatase (ALP) expression kit was received from Bioassay Systems (Hayward, CA). Hematoxylin and eosin-Y (H&E) for staining cell nuclei and cytoplasm, and Alcian blue for GAG staining were received from Sigma-Aldrich. All primary and secondary antibodies were received from Santa Cruz Biotechnology (Dallas, TX). All forward and reverse primers were synthesized and received from Integrated DNA Technologies (Coralville, IA).

### 2.2. Production of decellularized bovine cartilage microparticles

Full thickness articular cartilage samples, harvested from fetal or adult bovine femoral condyles, were decellularized as we described previously. Briefly, the articular cartilage samples were dissected with a scalpel into 5×5×2 mm pieces, the dissected pieces were frozen in liquid nitrogen and milled. The milled fragments were decellularized by immersion in 10 Mm Tris/1% triton solution for 24 h followed by sonication for 2 h at 55 kHz. Next, the sonicated fragments were immersed in nuclease solution consisting of 1 U/mL deoxyribonuclease and 1 U/mL ribonuclease in PBS for 72 h at 37°C to degrade DNA and RNA [22]. The decellularized fragments were washed 3X in PBS, centrifuged, the supernatant was discarded, and the solid was freeze-dried.

The freeze-dried fragments were further grinded (Hamilton Beach, Southern Pines, NC) and sorted for size by progressively passing through sieves ranging in size from 80 to 300 μm. First, the soft fragments were passed through 80 and 300 μm sieves to eliminate the <80 μm and >300 μm fragments. Next, the soft, decellularized cartilage microparticles (CMPs) were passed through a 90 μm sieve to collect a fraction with 40-110 μm size range, which is referred to as the 90 μm CMPs. Next, the >90 μm CMPs were passed through a 190 μm sieve to collect a fraction with 60-220 μm size range, which is referred to as the 190 μm CMPs. Then, the >190 μm CMPs were passed through a 250 μm sieve to collect a fraction with 60-300 μm size range, which is referred to as the 250 μm CMPs. The above process was repeated for the >250 μm fraction to sort the particles into the three 90, 190 and 250 μm fractions. Fetal and adult CMPs are hereafter referred to as fCMPs and aCMPs, respectively.

### 2.3. Characterization of the decellularized articular cartilage microparticles

#### Microparticle size distribution

The sieved CMPs were imaged with a light microscope to determine their size distribution. The captured 2D images were analyzed with ImageJ software (National Institutes of Health, Bethesda, MD) as we described previously [23]. After size analysis, the fetal and adult CMP fractions were immersed in liquid nitrogen and cut with a surgical blade to expose a freshly cut surface for morphological analysis. Next, the CMPs were coated with gold using a Denton Desk II sputter coater (Moorestown, NJ) at 20 mA for 75 s. The CMPs were imaged with a TESCAN VEGA3 SBU variable-pressure scanning electron microscope (SEM; Kohoutovice, Czech Republic) at an accelerating voltage of 8 keV.

#### Measurements of water content and mass loss

The equilibrium fractional water content of the dried, decellularized CMPs was measured by incubation in phosphate buffer saline (PBS) at 37°C as we described previously [24]. Briefly, after swelling in PBS, the CMPs were filtered, unbound water was removed with a filter paper, and weight of the samples was measured. After weighing, the samples were returned to fresh PBS solutions and incubated until the next time point. This process was repeated until equilibrium swelling was achieved. Equilibrium water content of the CMPs was calculated as the difference between the initial and swollen weights divided by the total weight of CMPs. Mass loss of the CMPs samples was measured at one time point after 8 weeks of incubation in PBS. After 8 weeks, the CMPs were filtered, freeze-dried, and weight of the samples was measured. Mass loss was defined as the difference between the initial and final weights of the dry CMPs divided by the initial dry weight.

### 2.4. Culture of MSCs on articular cartilage microparticles

MSCs (passage 3-5) were expanded in a high glucose DMEM medium supplemented with 10% FBS, 100 units/mL penicillin G, and 100 μg/mL streptomycin (basal medium, BM) as we previously described [18]. CMPs were sterilized by immersion in ethanol for 2 h followed by exposure to ultraviolet (UV) radiation for 30 min [25]. After washing, the sterilized adult or fetal CMPs were incubated in basal culture medium overnight to swell prior to cell seeding. The initial seeding density was calculated based on the average size of MSCs (18 μm average diameter [26]) and size and surface area of the CMPs. The initial cell seeding density was based on 1×10^6^ MSCs occupying 20% of the surface area of CMPs (20% initial confluency). For seeding 3×10^6^ cells, 0.22, 0.48 and 0.64 mg CMPs (adult or fetal) were required for average CMP sizes of 90, 190 and 250 μm, respectively. For 3D microcarrier-based cell culture, the specified amount of CMPs was added to the basal medium in ultra-low attachment T-75 tissue culture flasks followed by the addition of 3×10^6^ MSCs. The flasks were securely mounted on a rocker mixer and placed in a humidified 5% CO_2_ incubator at 37°C. A doubling time of 48 h is reported for MSCs cultured on natural ECMs [27]. Based on this doubling time, fresh CMPs were added to the cell culture medium every 48 h to limit confluency to 40% and prevent cell crowding on the surface of CMPs. The live/dead image analysis of the seeded MSCs confirmed the absence of cell crowding on the surface and in the pore volume of CMPs. MSCs cultured on 2D adherent culture flasks and MSCs cultured on 3D polystyrene microbeads (50 μm average size) were used as controls. At each time point (7, 14 and 21 days), the cultures were characterized with respect to cell number and viability, and mRNA expression of MSC markers.

### 2.5 Characterization of MSCs cultured on articular cartilage microparticles

#### Cell viability and growth

At each time point, 2 mL of 0.05% trypsin/ 0.53 mM EDTA was added to 1 mL of the cell culture suspension and incubated for 15 min under shaking to detach MSCs from the CMPs. Next, the suspension was transferred to a 40 μm nylon cell strainer fixed on a 50 mL Falcon tube and washed with DMEM using an insulin syringe. The filtrate was centrifuged at 400×g for 5 min as we described previously [28], and the separated cells were counted with a hemocytometer [28]. For cell viability, the CMPs were stained with 1 μg/mL live/dead cAM/EthD and imaged with an inverted fluorescent microscope (Nikon Eclipse Ti-e, Nikon, Melville, NY), as we described previously [24].

#### Analysis of mRNA expression

The MSCs cultured on CMPs were characterized phenotypically by mRNA upregulation of CD105, CD166 and CD44 markers, and downregulation of CD45 and CD34 markers [29]. Further, the MSCs cultured on CMPs were tested for differentiation to the chondrogenic lineage by measuring mRNA expression of chondrogenic markers SOX-9, Collagen I (Col I), Collagen II (Col II), and aggrecan (AGC) [17, 18]. At each time point, MSCs were separated from the CMPs as described above and the total RNA of the homogenized cell suspension was isolated using TRIzol as we described previously [23]. The genomic DNA was removed using deoxyribonuclease I (Invitrogen) as previously described [16]. 250 ng of the extracted RNA was converted to cDNA using Promega reverse transcription system (Madison, WI). The cDNA was amplified with Eppendorf SYBR green RealMasterMix (Hamburg, Germany) using a Bio-Rad CXF96 real-time quantitative polymerase chain reaction system (rt-qPCR; Hercules, CA) and the appropriate gene-specific primers as descried [17]. The primer sequences, listed in Table 1, were designed and selected using Primer3 web-based software as described [16]. The expressions were normalized against GAPDH reference gene and fold changes were compared based on ΔΔct method to those in the same group at day zero as previously described [30].

**Table 1.**
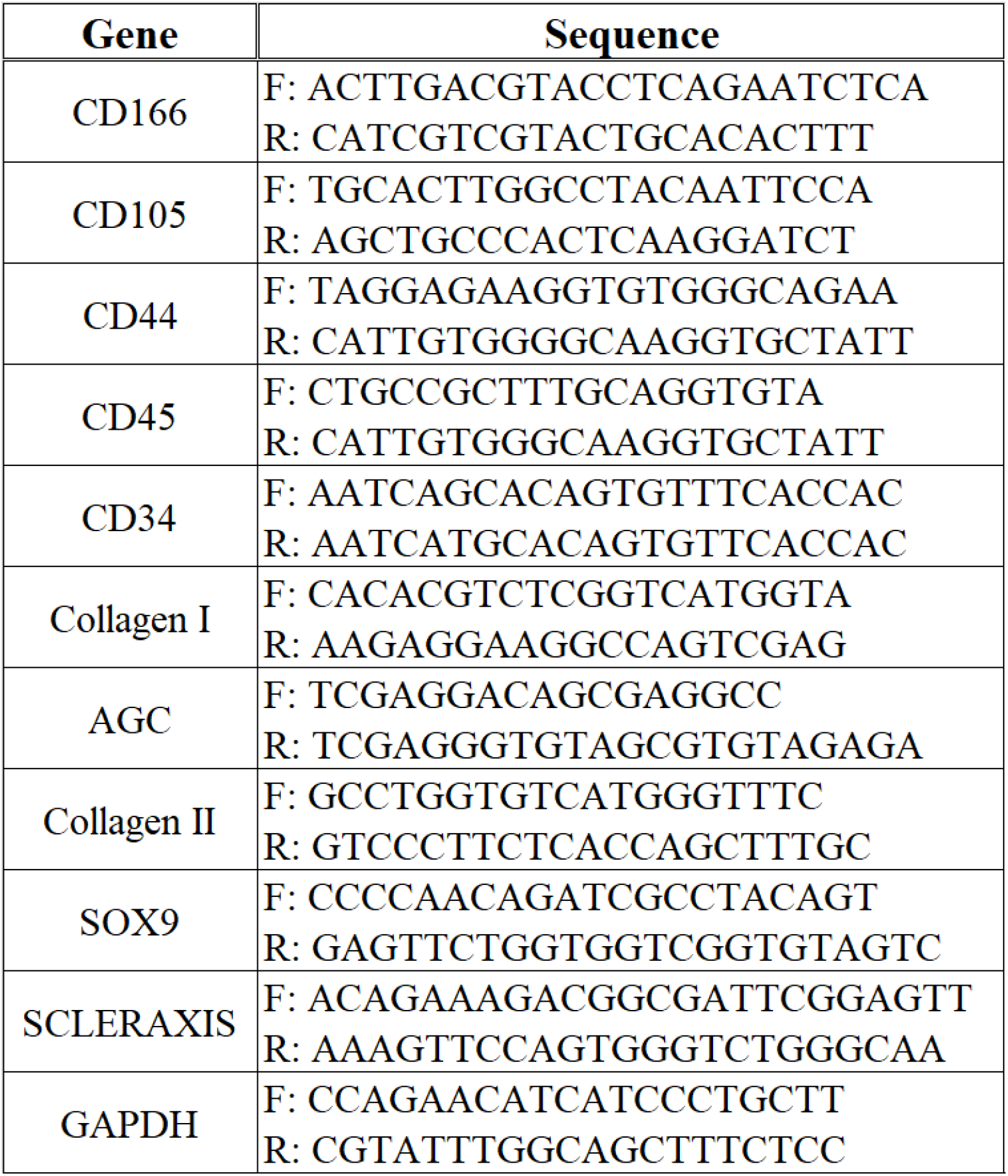
Forward and Reverse sequences for PCR primers.

### 2.6. Encapsulation of CMP-MSCs in alginate as an injectable hydrogel

The following approach was used to generate CMP-MSC encapsulated alginate gels as an injectable cell carrier for articular cartilage regeneration [31]. The alginate (3 g sodium alginate to 100 mL PBS) and CaCl_2_ (1 g CaCl_2_ to 100 mL PBS) solutions [25] were sterilized by filtration. A suspension of adult or fetal CMP-MSCs in culture medium, prepared as described in section 2.4, was transferred to a sterile 15 mL Falcon tube, centrifuged, medium was removed, and the alginate solution was added to the CMP-MSCs followed by mixing with a pre-sterilized glass rod. Next, the surface of a pre-sterilized, disk-shape, Teflon mold with effective diameter of 2 cm and height of 1.5 mm was sprayed with the 1% CaCl_2_ solution using an insulin syringe. Then, the suspension of CMP-MSCs in sodium alginate was transferred to the mold and the exposed surface area of the mold was sprayed with the CaCl_2_ solution to crosslink the suspension. The gelation time was determined visually by placing a drop of the suspension on a glass ruler, spraying the drop with the CaCl_2_ solution, tilting the ruler by 45 degrees, and recording the time for the drop to stop flowing. The gelation time of the CMP-MSC encapsulated in alginate ranged from 1-4 min. After crosslinking, the cell-encapsulated gels were transferred to a petri dish and cultured in chondrogenic medium for up to 8 weeks. MSCs encapsulated directly in the alginate gel, without CMPs, at a density of 1×10^5^ cells/mL were used as the control group. aCMP-MSCs and fCMP-MSCs encapsulated in alginate gels are hereafter referred to as aCMP-MSCs/alg and fCMP-MSCs/alg, respectively.

### 2.7. Formation of CMP-MSC monolayer and bilayer cell sheets

The following approach was used to generate CMP-MSC cell sheets as a preformed cell carrier for articular cartilage regeneration. A suspension of adult or fetal CMP-MSCs (approximately 50% cell confluency) in culture medium, prepared as described in section 2.4, was transferred to a sterile Teflon mold (1.55 mm depth and 2 cm in diameter), the mold was placed in a sterile petri dish, and the assembly was incubated in a humidified 5% CO_2_ incubator at 37°C for 48 h. Gravitational settling of the CMP-MSCs on the bottom surface of the mold followed by ECM secretion led to the formation of adult or fetal CMP-MSCs monolayer cell sheets, hereafter referred to as aCMP-MSCs/ml or fCMP-MSCs/ml cell sheet, respectively. The CMP-MSC loading in the culture medium was varied to form cell sheets with 0.75 mm thickness. The following approach was used to produce bilayer cell sheets. After formation of aCMP-MSC/ml, a suspension of fCMP-MSCs was transferred to the mold with aCMP-MSC/ml cell sheet, and the assembly was incubated for 48 h to gravitational settle fCMP-MSCs on top of aCMP-MSCs/ml cell sheet followed by ECM secretion to form a bilayer cell sheet, hereafter referred to as faCMP-MSCs/bl cell sheet. The CMP-MSC loading in the culture medium was adjusted to form faCMP-MSCs/bl cell sheets with 1.5 mm thickness. After cell sheet formation, the medium was replaced with chondrogenic medium and the cell sheets were cultured for up to 8 weeks. The chondrogenic medium consisted of DMEM (4.5 g/mL glucose, 50 μg/ mL L-proline, 50 μg/mL ascorbic acid, 0.1 mM sodium pyruvate, 1% v/v insulin-transferrin-selenium premix) supplemented with 10 ng/mL TGF-β1 [16]. MSC pellets formed directly by centrifugation were used as the control group [32].

### 2.8. Analysis of monolayer, bilayer, or alginate encapsulated CMP-MSCs

At each time point, adult or fetal CMP-MSCs/alg, CMP-MSCs/ml or CMP-MSCs/bl were assessed with respect to compressive modulus, cellularity, and the expression of chondrogenic markers Sox-9, Collagen II (Col II) and aggrecan (AGC), the superficial zone marker SZP, and calcified zone markers collagen X (Col X) and alkaline phosphatase (ALP). Cell viability was assessed by imaging with live/dead cell assay. The mono/bi-layer sheets were incubated with cAM/EthD live/dead stains, as we described previously [16], and the stained samples were imaged using an Eclipse Ti-E inverted fluorescent microscope. For cell imaging, CMP-MSCs/alg gels were fixed with 4% paraformaldehyde for 3 h, permeabilized using PBS containing 0.1% Triton X-100 for 5 min, and incubated with Alexa 488 phalloidin (1:200 dilution) and DAPI (1:5000 dilution) to stain actin filaments of the cell cytoskeleton and cell nuclei, respectively, as previously described [33]. The stained gels were imaged with the Eclipse inverted fluorescent microscope. The mRNA expression of chondrogenic markers of MSCs in the samples were measured as described in section 2.5.

### 2.9. Histological Analysis

After 21 days of culture, the expression of GAG and mineralized deposits in aCMP-MSCs/ml, fCMP-MSCs/ml, or faCMP-MSCs/bl was measured histologically as we previously described [16]. Briefly, the samples were fixed in formalin, embedded in paraffin, and cryo-sectioned to a thickness of 10 μm. The sections were divided into three groups with the first group stained with H&E to ascertain morphology of the encapsulated cells, the second group stained with Alcian blue to image GAG accumulation, and the third group stained with Alizarin red to image mineral deposit. The stained sections were imaged with a Nikon Optiphot Epi-fluorescent microscope.

### 2.10. Compressive modulus of monolayer, bilayer, or alginate encapsulated CMP-MSCs

At each time point, adult or fetal CMP-MSCs/alg, CMP-MSCs/ml, or CMP-MSCs/bl were loaded on the Peltier plate of an AR 2000 rheometer (TA Instruments, New Castle, DE, USA) and subjected to a uniaxial compressive strain as we previously described [34]. A strain sweep was performed from 0.01% to 500% strain at 10 Hz to determine the yield strain. Similarly, a frequency sweep was performed from 0.01 to 100 Hz at 0.2% strain to determine the crossover frequency. A sinusoidal shear strain with a frequency above the crossover frequency and a strain amplitude below the yield strain was exerted on the sample and the storage (G’) and loss moduli (G”) were recorded with time. The slope of the linear fit to the stress-strain curve for strains of <10% was taken as the compressive modulus of the samples.

### 2.11. Statistical Analysis

All experiments were done in triplicate and quantitative data was expressed as means + standard deviation. Significant differences between groups were evaluated using a two-way ANOVA with replication test and two-tailed Student’s t-test. A value of p < 0.05 was considered statistically significant.

## 3. Results and discussion

Figures 2a through 2c show size distribution of aCMPs with average sizes of 90 μm, 190 μm and 250 μm, respectively; Figures 2d through 2f show size distribution of fCMPs with average sizes of 90 μm, 190 μm and 250 μm; the insets in Figures 2a-f show the corresponding microscope images of the CMPs. For fetal as well as adult CMPs and for the three sizes, the fraction corresponding to the average size was between 70-90% of the distribution. The fraction of particles with size greater than the average was higher for fCMPs compared to aCMPs, which was attributed to the softer texture of fCMPs and their passage through the sieve with a slight pressure. The harder texture of aCMPs resulted in a smaller particle size distribution compared to the fetal. The CMPs had irregular, non-spherical shapes and there was no difference in the shape of fetal and adult CMPs.

**Figure 2.**
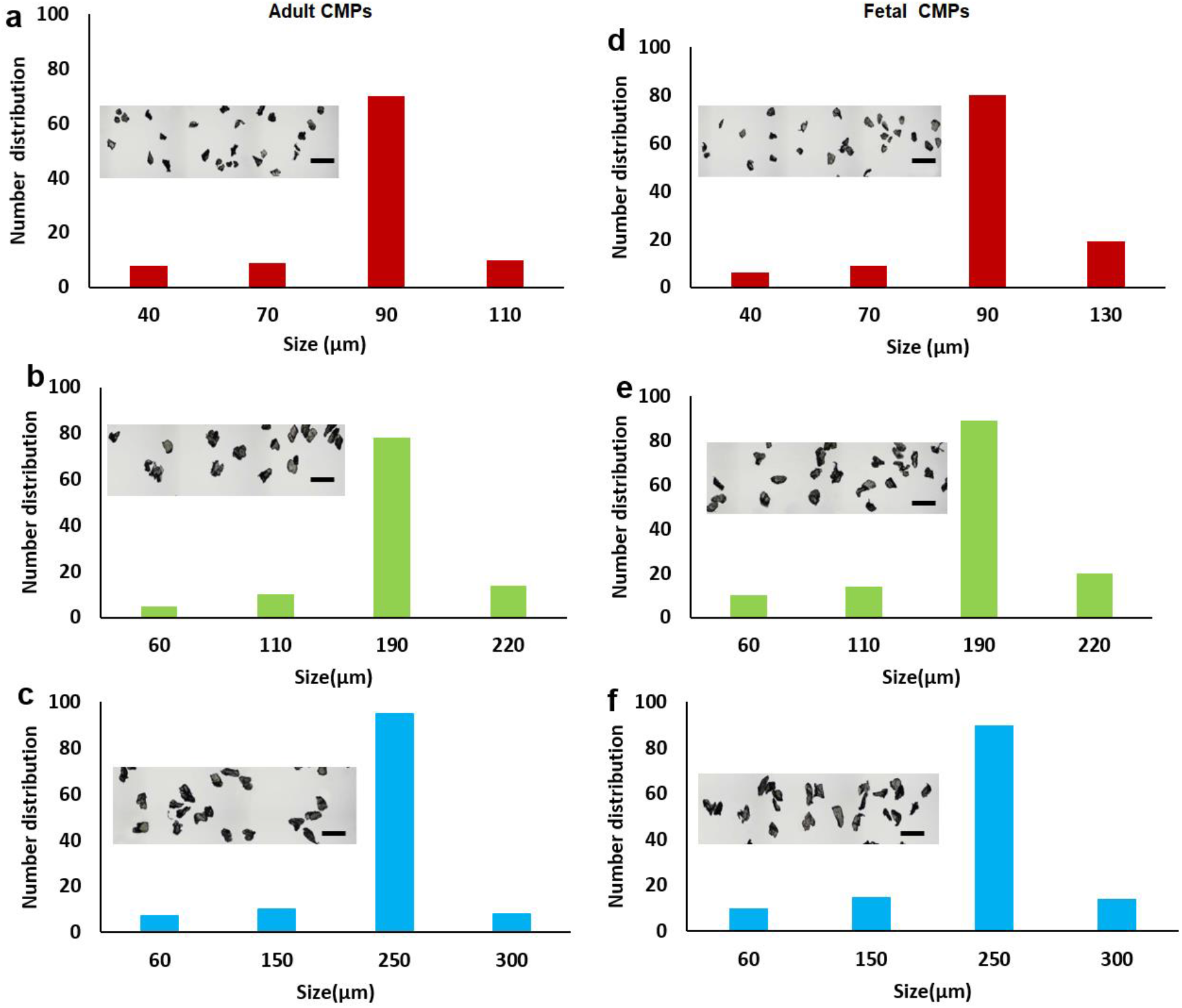
The size distribution of adult (a-c) and fetal (d-f) CMPs. Distributions a, b and c correspond to adult CMPs with average sizes of 90, 190 and 250 μm, respectively; Distributions d, e and f correspond to fetal CMPs with average sizes of 90, 190 and 250 μm; the insets in the distributions are SEM images of the CMPs.

The percent equilibrium water content of fCMPs with average particle sizes of 90 μm, 190 μm and 250 μm was 18.6±0.9%, 19.6±0.9% and 20.0±1.5%, respectively, and that of aCMPs was 15.3±1.5%, 16.5±0.6% and 18.0±0.8%. The water content of fCMPs was not affected by their particle size whereas that of aCMPs increased slightly with increasing size. For all particle sizes, the water content of fCMPs was higher than aCMPs. The mass loss of fCMPs with average particle sizes of 90 μm, 190 μm and 250 μm after 8 weeks incubation in PBS was 4.4±0.2%, 5.2±0.2% and 6.5±0.1%, respectively; that of aCMPs was 3.7±0.3%, 4.3±1.3% and 5.8±0.5%. The mass loss of fetal and adult CMPs increased with increasing particle size. For a given particle size, the mass loss of aCMPs was slightly lower than fCMPs. The mass loss data indicated that the fetal and adult CMPs are stable for up to 8 weeks in the absence of enzymes with negligible hydrolytic degradation.

Two culture methods were used for expansion of MSCs on CMPs. In the first method, all CMPs were suspended in basal culture medium initially along with MSCs at time zero. In the second method, all MSCs were added to the culture medium initially at time zero but the CMPs were added gradually every two days starting. The second method was more efficient as the surface area for cell adhesion and growth was increased gradually without cell confluency falling below 20%, which improved cell-cell interaction between the adhered MSCs. The second method allowed continuous expansion of MSCs by intermittent adding of CMPs to the culture medium with incubation time. Figures 3a and 3b compare the cell content of fetal and adult CMPs, respectively, with average sizes of 90 μm, 190 μm and 250 μm with incubation time against MSCs grown on 3D polystyrene microparticles (PSMP). For all time points, the cell content of the CMP groups was higher than the control and increased with incubation time. For all time points, the fetal or adult CMPs with 250 μm average size had highest cell content compared to other CMP sizes. The lower cell content of the 90 μm CMPs compared to 250 μm CMPs was attributed to higher inter-particle cell transfer with each successive CMP addition to the culture medium. Inter-particle transfer requires cell detachment from one particle, migration through the medium, and re-attachment to another particle with lower cell content, which increased the lag time between cell divisions. Figure 3c compares the cell content of fetal and adult CMPs as a function of average particle size with incubation time. For a given time and particle size, the cell content of fCMPs was slightly higher than aCMPs but the difference was not statistically significant.

**Figure 3.**
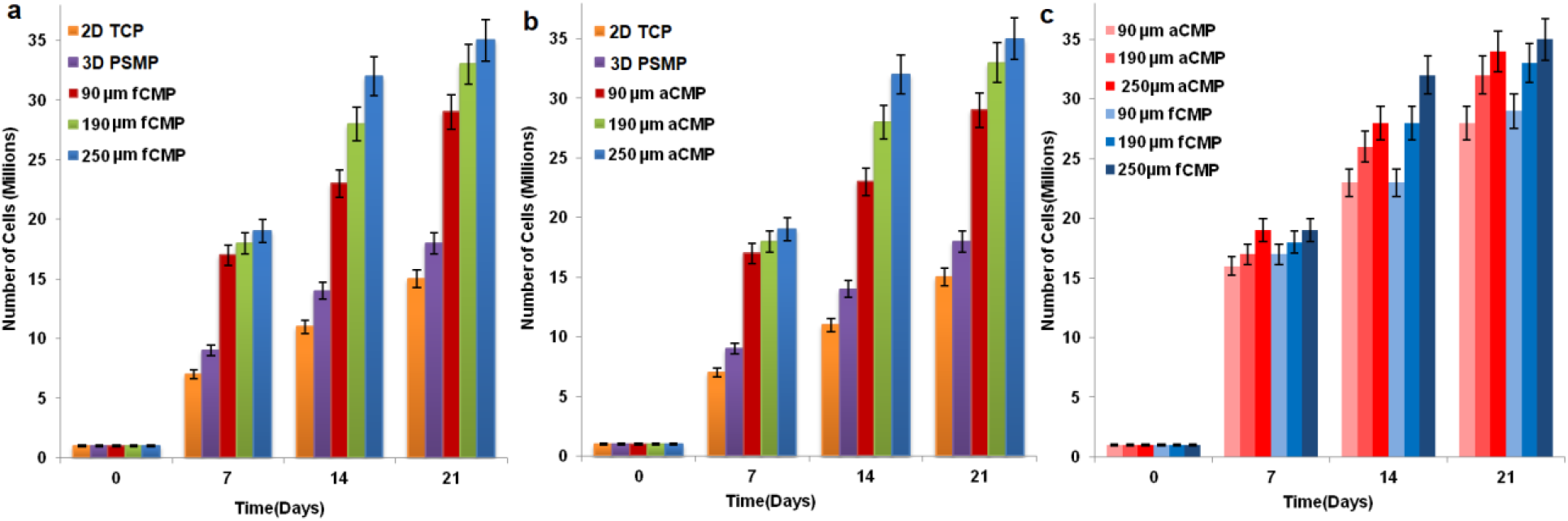
Expansion of MSCs on fetal (a) and adult (b) CMPs with incubation time in basal medium in a tissue culture bioreactor for CMP particle sizes of 90 μm (red), 190 μm (green) and 250 μm (blue); MSCs grown on 2D tissue culture plates (orange) and 3D polystyrene beads (purple) were used as controls; (c) comparison of MSC growth on fetal (blue) and adult (red) CMPs as a function of incubation time in basal medium for CMP particle sizes of 90 μm (light), 190 μm (medium) and 250 μm (dark).

The live (green) and dead (red) fluorescent images of a randomly selected particle in fetal or adult CMP samples are shown in Figure 4 as a function of incubation time for different average sizes. Based on fluorescent images, the MSCs penetrated the pore structure of CMP particles. The intensity of green fluorescence from the CMPs increased with incubation time for fetal as well as adult CMPs and for all particle sizes. After 21 days of incubation, the fluorescent intensity of fCMPs was slightly higher than aCMPs for all particle sizes but the difference was not statistically significant. The fluorescent images of CMP-MSCs showed >95% cell viability.

**Figure 4.**
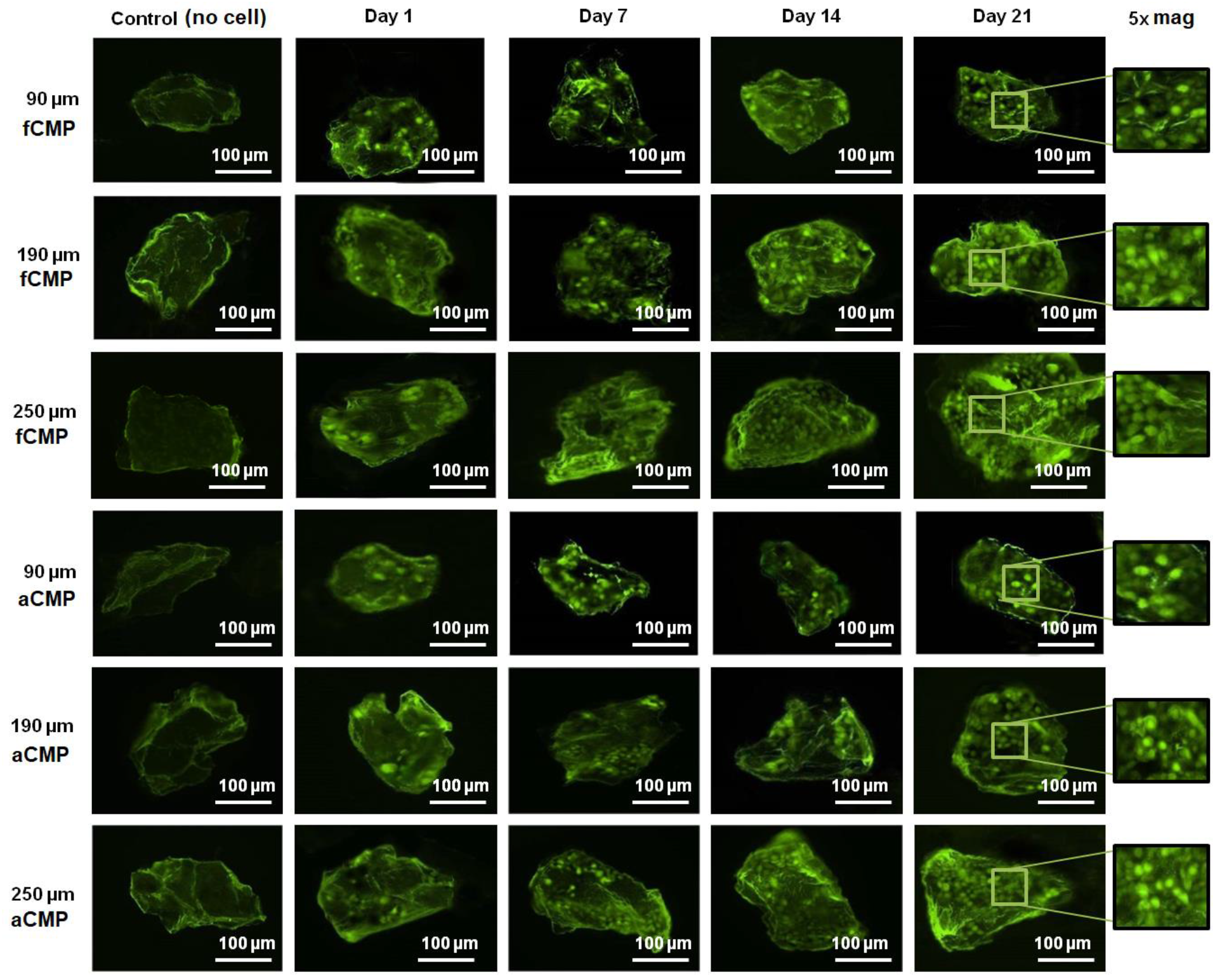
Live (green) and dead (red) fluorescent images of randomly selected microparticles from fetal and adult CMP-MSCs with incubation time in basal medium for CMP particle sizes of 90, 190 and 250 μm. The control group (most left column) is the image of a microparticle without incubation with MSCs. The images in the last column on the right are 5x magnification of the 21 days images. The scale bar in the images is 100 μm.

According to previous reports, the expression of CD105 and CD44 in human MSCs harvested from the bone marrow is upregulated whereas the expression of CD45 and CD34 is down regulated [35, 36]. Further, the as-received MSCs had high expression of CD105, CD166, CD29, and CD44 markers and low expression of CD14, CD34 and CD45. The expression of CD105, CD166 and CD44, CD45 and CD34 markers for the MSCs expanded on fetal/adult CMPs are shown in Figure 5. According to this figure, the expression of CD105, CD166 and CD44 markers of MSCs expanded on fetal/adult CMPs cultured in basal medium increased with incubation time whereas there was absence of CD45 and CD34 expression. There was no difference between the marker expression of CMP-MSCs and 2D cultured MSCs; there was also no difference between the marker expression of adult and fetal CMP-MSCs. It is possible for CMPs to prematurely induce differentiation of MSCs during the expansion phase. Therefore, in addition to MSC markers, the expression of chondrogenic and osteogenic markers of CMP-MSCs were measured. These included Sox-9 as the master regulator of chondrogenesis, Col II as the chondrogenic marker, and Col I as the osteogenic marker (Figure 5) [37]. According to Figure 5, the expression of Sox-9, Col II and Col I of CMP-MSCs decreased with incubation time in basal medium. Further, there was no difference in the expression of chondrogenic markers between adult and fetal CMP-MSCs. These results demonstrate that MSCs can be expanded on fetal or adult CMPs in basal medium without premature differentiation to the chondrogenic or osteogenic lineages.

**Figure 5.**
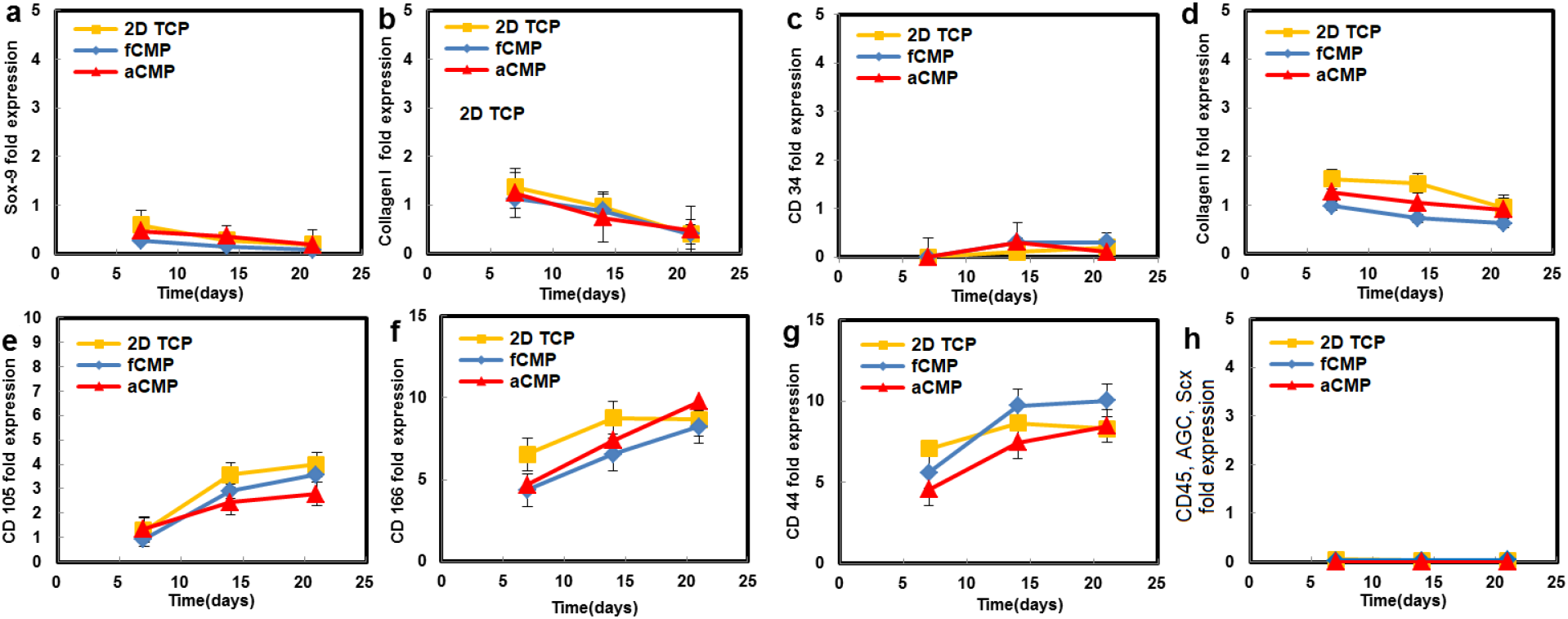
mRNA expression of Sox-9 (a), Col I (b), CD34 (c), Col II (d), CD105 (e), CD166 (f), Cd44 (g), and CD45, AGC, Scx (h) markers of MSCs cultured on adult (red) and fetal (blue) CMPs as a function of incubation time in basal medium for 21 days. The control group (yellow) is MSCs cultured on 2D tissue culture plates.

Injectable and preformed approaches were used to generate cell carriers for articular cartilage regeneration. Figure 6a shows the effect of CMP size on gelation time of the alginate in the injectable approach for different concentrations of CaCl_2_. The pure alginate solution had the fastest gelation time compared to alginate solutions with CMP-MSCs for all CaCl_2_ concentrations. For a given CMP size, gelation time decreased almost linearly with increasing CaCl_2_ concentration. For a given CaCl_2_ concentration, gelation time increased with increasing CMP particle size. Figure 6b shows the effect of CMP loading on gelation time of the alginate solution for different concentrations of CaCl_2_. For a given CaCl_2_ concentration, the gelation time decreased with increasing CMP loading from 30% to 70% by volume. In general, the gelation times were in the range of 1-4 min, which were within the clinically acceptable range for injectable cell-encapsulated hydrogels [38].

**Figure 6.**
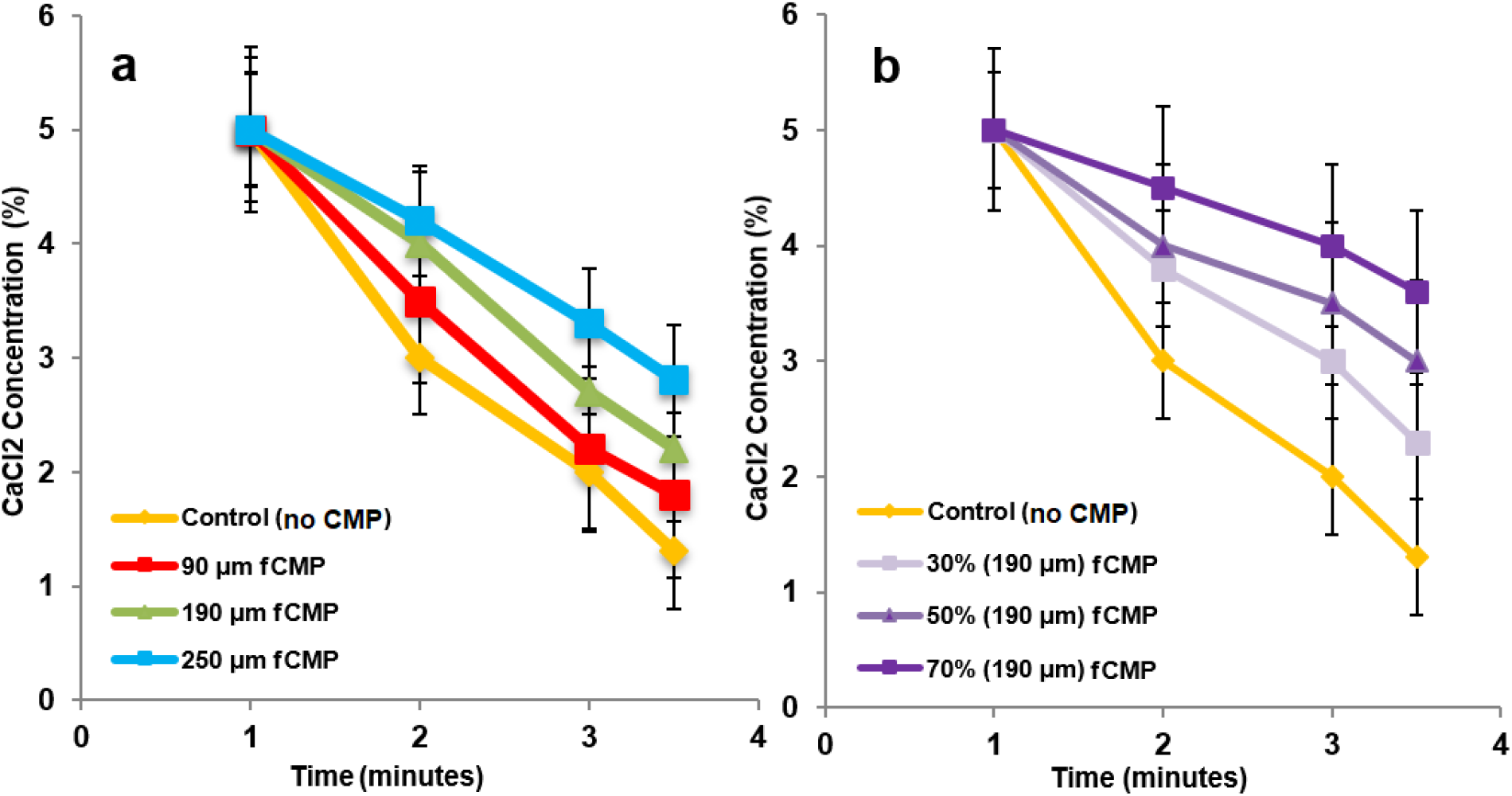
(a) Gelation time of alginate gels as a function of CaCl_2_ concentration containing 50% (by alginate weight) CMPs with particle sizes of 90 μm (red), 190 μm (green), and 250 μm (blue); (b) Gelation time of alginate gels as a function of CaCl_2_ concentration containing 30% (very light purple), 50% (light purple) and 70% (purple) CMPs with average size of 190 μm. The control group in (a,b) is alginate gel without CMPs.

The graphs in exhibit A of Figure 7 compare mRNA expression of chondrogenic markers for adult or fetal CMP-MSCs/alg hydrogels as a function of incubation time in chondrogenic medium. The control group in exhibit A was MSCs encapsulated directly, without CMPs, in alginate. Chondrogenic markers included Sox-9 [37], SZP as the superficial zone marker [39], Col II and AGC as the middle zone markers [16], and Col X and ALP as calcified zone markers [40]. For all markers and incubation times, adult or fetal CMP-MSCs showed higher expressions compared to directly encapsulated MSCs in alginate. There was no difference between the expressions of Sox-9, AGC, Col X and ALP of aCMP-MSCs/alg and fCMP-MSCs/alg whereas SZP was higher in aCMP-MSCs/alg and Col II was lower. As the culture medium was not supplemented with zone-specific growth factors, we did not expect a significant difference in the expression of zone-specific markers between aCMP-MSCs and fCMP-MSCs, consistent with our previous results that matrix composition and zone-specific growth factors work synergistically to enhance MSC differentiation to zone-specific chondrogenic phenotypes [16]. According to the data in exhibit A of Figure 7, adult or fetal CMP-MSCs enhanced chondrogenic differentiation of MSCs in the alginate hydrogel, as compared to the commonly used method of direct encapsulation of MSCs in the gel [41].

**Figure 7.**
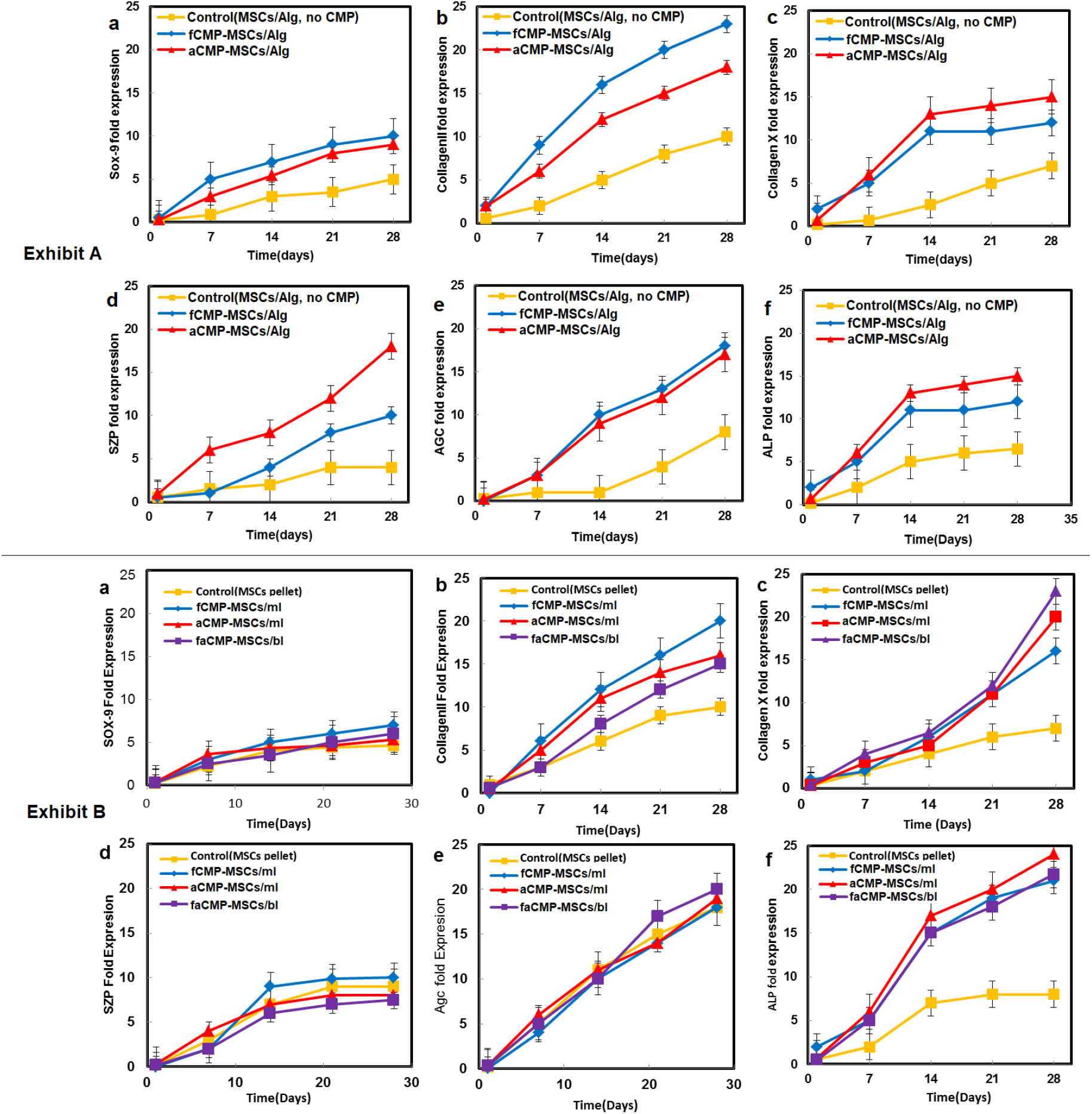
(Exhibit A) mRNA expression of chondrogenic markers Sox-9 (a), Col II (b), Col X (c), SZP (d), AGC (e), and ALP (f) with incubation time for fCMP-MSCs (blue) or aCMP-MSCs (red) encapsulated in injectable alginate hydrogels and incubated in chondrogenic medium for 28 days; the control group (yellow) in Exhibit A is MSCs (without CMP) encapsulated in alginate hydrogel and incubated in chondrogenic medium; (Exhibit B) mRNA expression of chondrogenic markers Sox-9 (a), Col II (b), Col X (c), SZP (d), AGC (e), and ALP (f) with incubation time for CMP-MSCs preformed cell sheets and incubated in chondrogenic medium for 28 days; groups include monolayer fCMP-MSCs (blue), monolayer aCMP-MSCs (red), and bilayer faCMP-MSCs (purple) cell sheets; the control group (yellow) in Exhibit B is the MSC pellet cultured in chondrogenic medium.

In the preformed approach, mono- or bilayer cell sheets were generated by sequential gravitational settling of aCMP-MSCs or fCMP-MSCs on the culture plate followed by cell fusion with incubation. Microscopic images of fCMP-MSCs, aCMP-MSCs and faCMP-MSCs cell sheets stained with blue, red, and purple dyes, respectively, are shown in Exhibit A of Figure 8. The dye-stained images in Exhibit A show progressive fusion and disappearance of the interface between CMP-MSCs with incubation time from 2 to 6 days for monolayer and bilayer cell sheets. The Exhibit B in Figure 8 compares the elastic modulus of CMP-MSCs cell sheets with MSC pellets (without CMPs) as a function of incubation time in chondrogenic medium. The elastic moduli of CMP-MSC cell sheets and the MSC pellet increased steadily with incubation time. For a given incubation time, the elastic moduli of CMP-MSCs cell sheets were higher than the MSC pellet, which was attributed to enhanced chondrogenic differentiation and ECM secretion of ECM-MSCs. Further, the elastic moduli of fCMP-MSCs monolayer and faCMP-MSCs bilayer were higher than aCMP-MSCs monolayer after 8 weeks of incubation.

**Figure 8.**
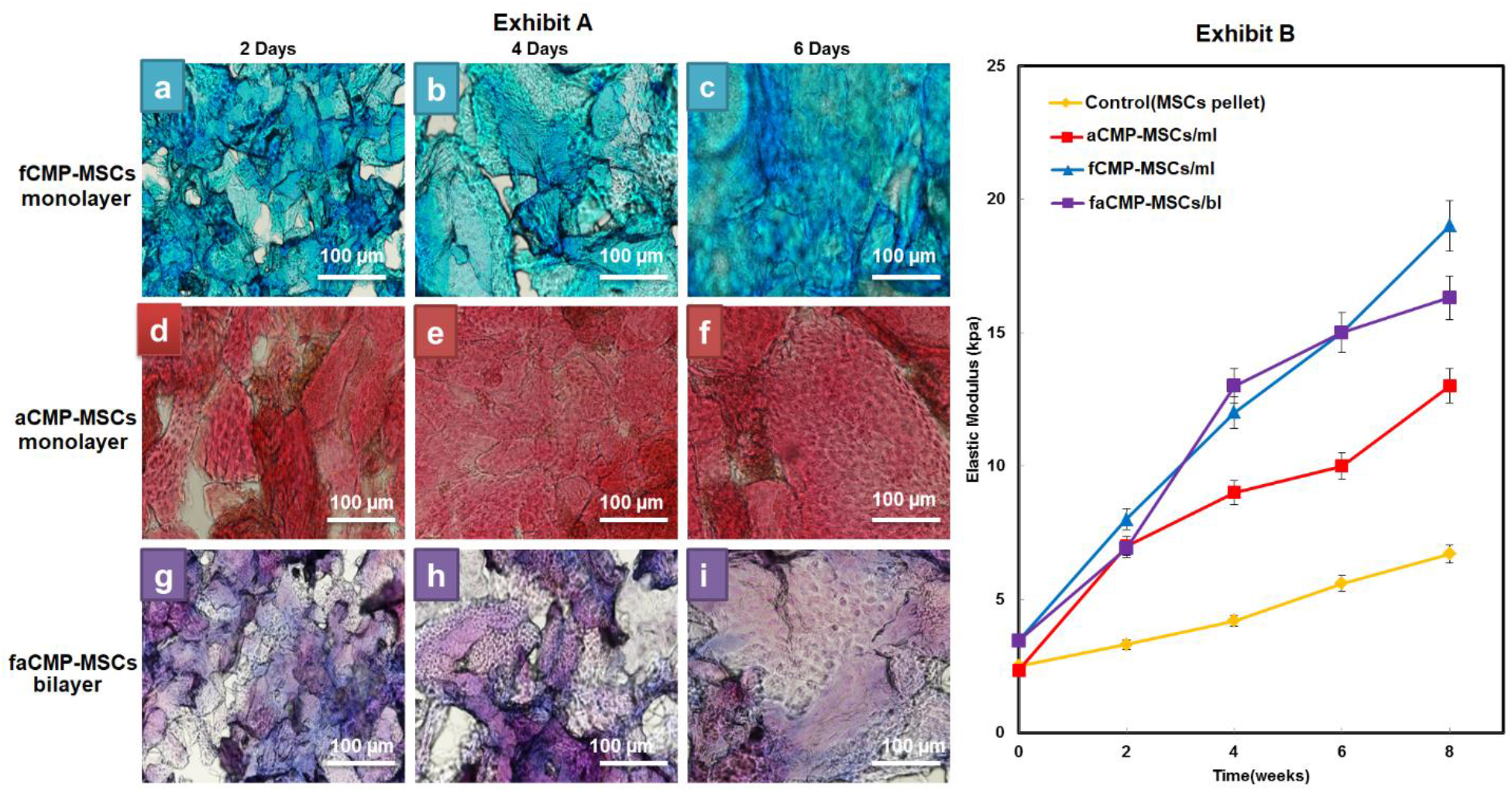
(Exhibit A) images of monolayer fCMP-MSCs (blue), monolayer aCMP-MSCs (red), and bilayer faCMP-MSCs (purple) cell sheets after 2, 4, and 6 days incubation in chondrogenic medium stained with blue, red, and purple dyes, respectively; the scale bar in the images is 100 μm; (Exhibit B) elastic modulus of fCMP-MSCs (blue), aCMP-MSCs (red), and bilayer faCMP-MSCs (purple) cell sheets with incubation time in chondrogenic medium for 6 days; the control group in Exhibit B is the MSC pellet cultured in chondrogenic medium.

The graphs in exhibit B of Figure 7 compare mRNA expression of chondrogenic markers for aCMP-MSCs/ml, fCMP-MSCs/ml, and faCMP-MSCs/bl as a function of incubation time in chondrogenic medium. The control group in exhibit B is the commonly used 3D pellet culture [32]. For Sox-9, SZP and AGC markers, the expressions for CMP-MSCs monolayers and bilayers were relatively close to the pellet culture whereas for Col II, Col X and ALP makers, the expressions for CMP-MSCs were higher than the pellet culture. There was not a difference in marker expressions between the adult and fetal CMP-MSCs cell sheets, except for Col II of fCMP-MSCs/ml which was slightly higher than faCMP-MSCs/bl. The data in Figures 5–7 show that the MSCs expanded CMPs could potentially be injected or implanted in an articular cartilage defect without the need to detach and separate MSCs from CMPs. Further, the experimental results indicate that the adult or fetal CMPs, as a biomimetic microcarrier, enhance chondrogenic differentiation and maturation of MSCs.

Figure 9 shows the compressive moduli of adult and fetal CMP-MSCs/alg hydrogels as a function of CMP content with incubation time in chondrogenic medium; the moduli of aCMP-MSCs/alg with average CMP sizes of 90, 190 and 250 μm are shown in Figures 9a-c, respectively; and the moduli of fCMP-MSCs with sizes of 90, 190 and 250 μm are shown in Figures 9d-f. The compressive moduli of all samples including the control group (MSCs directly encapsulated in alginate gel) increased with incubation time. All adult or fetal CMP-MSC groups reached their highest compressive modulus after 8 weeks of incubation. For a given incubation time, the compressive moduli of adult or fetal CMP-MSCs/alg groups were higher than that of the control (yellow curve) irrespective of CMP percent or CMP size. The moduli of adult of fetal CMP-MSCs/alg groups with 50% and 70% CMP were higher than the 30% CMP group for all CMP sizes and incubation times. The moduli of adult or fetal CMP-MSCs/alg groups with 190 μm and 250 μm particle sizes were higher than the 90 μm size group for all CMP loadings and incubation times. The adult CMP-MSCs/alg with 250 μm CMP size with 50% CMP loading after 8 weeks incubation had highest compressive modulus of 238±10 kPa followed by the group with 190 μm CMP size and 50% CMP loading at 218±10 kPa. The fetal CMP-MSCs/alg groups with 250 μm and 190 μm CMP sizes and 50% CMP loading after 8 weeks incubation had highest modulus of 197±20 kPa. Overall, the adult CMP-MSCs/alg with 250 μm CMP size, 50% CMP loading, and 8 weeks of incubation had highest compressive modulus. The higher modulus of adult or fetal CMP-MSCs/alg groups with CMP sizes of 250 and 190 μm is attributed to the higher number of cells per CMP, resulting in higher extent of cell-cell contact. This is consistent with previous reports that cell-cell contact enhances chondrogenic differentiation of MSCs.

**Figure 9.**
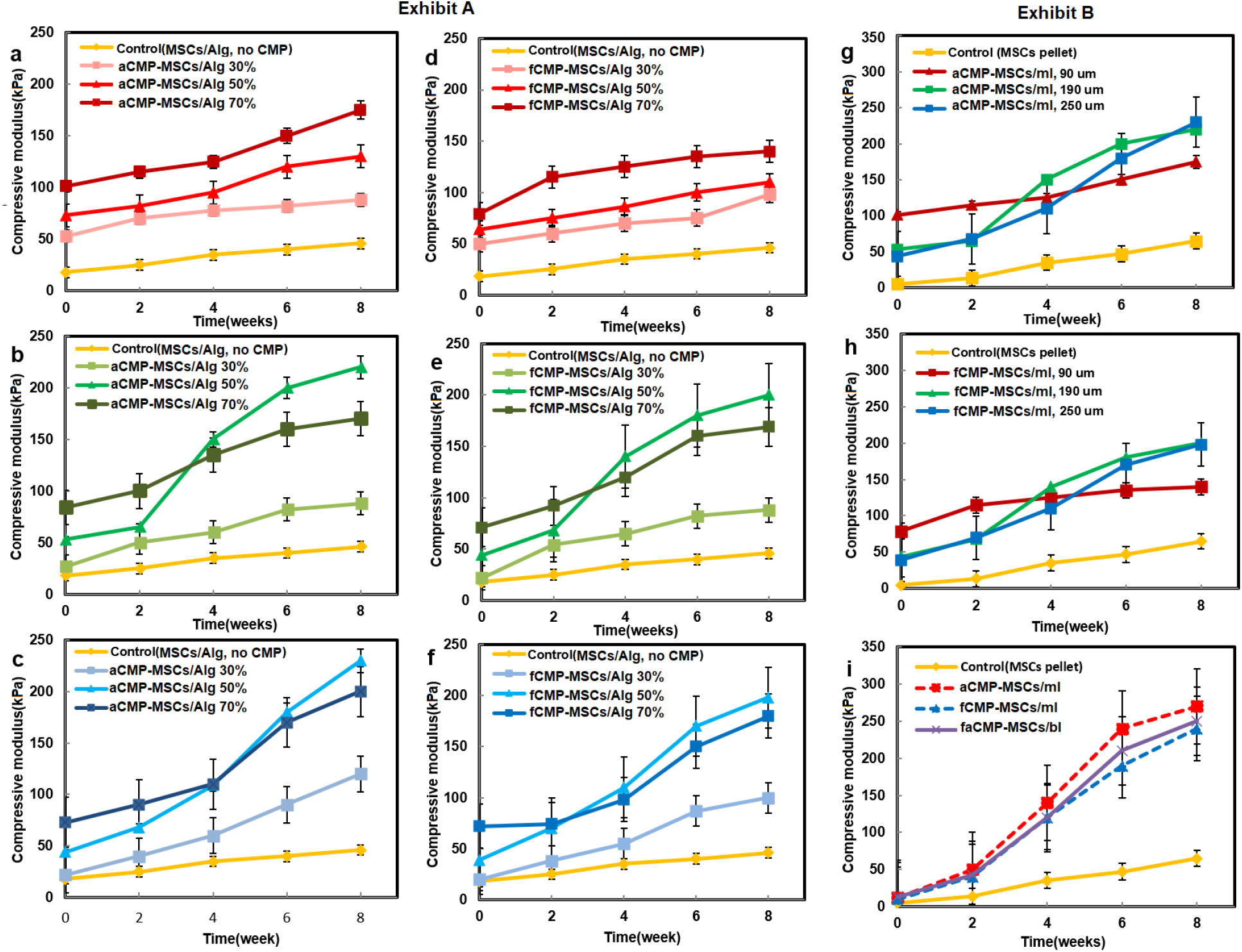
(Exhibit A) compressive modulus of aCMP-MSCs/Alg (left column) and fCMP-MSCs/Alg (right column) with incubation time in chondrogenic medium as a function of percent CMP and average CMP size; CMP percentages in Exhibit A were 30% (light shade), 50% (medium shade), and 70% (dark shade); Figures a, b, and c in Exhibit A are for aCMP-MSCs/alg with average CMP sizes of 90 μm (a), 190 μm (b) and 250 μm (c), respectively, whereas Figures d, e, and f are for fCMP-MSCs/alg; the control group (yellow) in Exhibit A is MSCs (without CMP) encapsulated in alginate hydrogel and incubated in chondrogenic medium; (Exhibit B) compressive modulus of aCMP-MSCs/ml (g) and fCMP-MSCs/ml (h) with incubation time in chondrogenic medium as a function of average CMP sizes of 90 μm (red), 190 μm (green) and 250 μm (blue); (i) comparison of compressive moduli of aCMP-MSCs/ml (dash red), fCMP-MSCs/ml (dash blue), and faCMP-MSCs/bl (solid purple) cell sheets as a function of incubation time in chondrogenic medium; the control group (yellow) in Exhibit B is the MSC pellet cultured in chondrogenic medium.

Figures 9g-i show the compressive moduli of mono- and bilayer cell sheets as a function of CMP size with incubation time in chondrogenic medium; the moduli of adult and fetal CMP-MSCs/ml cell sheets as a function of CMP size is shown in graphs g and h, respectively; the moduli of adult and fetal CMP-MSCs/ml and faCMP-MSCs/bl are compared in graph i. The control group in Figures g-i was MSCs in pellet culture [38]. The moduli of all CMP-MSCs cell sheets increased continuously with incubation time. The moduli of CMP-MSCs cell sheets were higher than the control group (MSC pellet) for all incubation times. The moduli of adult or fetal CMP-MSCs cell sheets with 250 and 190 μm CMP sizes were higher than the 90 μm CMP size (Figures 8g,h) after 8 weeks of incubation, which was attributed to higher extent of cell-cell interaction in larger CMPs [42]. The moduli of aCMP-MSCs/ml and fCMP-MSCs/ml, produced from CMP-MSCs with equal parts of 90, 190 and 250 μm CMPs, and faCMP-MSCs/bl were higher than the MSC pellet group for all time points. However, after 8 weeks of incubation, there was no difference between the moduli of aCMP-MSCs/ml, fCMP-MSCs/ml, and faCMP-MSCs/bl groups. The moduli of CMP-MSCs monolayer and bilayer cell sheets were in the range of 250±30 kPa which was higher than that of the MSC pellet group at 70±5 kPa. Overall, after 8 weeks of incubation, the compressive moduli of adult or fetal CMP-MSCs/ml cell sheets (250±30 kPa) were higher than the moduli of CMP-MSCs/alg (238±10 kPa) hydrogels.

Exhibit A in Figure 10 shows cAM/EthD stained images of live/dead cells in CMP-MSCs/alg and CMP-MSCs cell sheets, respectively; exhibit B shows the corresponding phalloidin/DAPI stained images of the shape and position of the viable cells. The first and second rows in exhibits A and B of Figure 10 correspond to fetal and adult CMPs, respectively, whereas the third row is for fa/CMP-MSCs/bl cell sheet. The injectable CMP-MSCs/alg hydrogels in Figure 10 had 50% CMP loading with average size of 190 μm. The cell sheets in Figure 10 were generated from CMP-MSCs with equal parts of 90, 190 and 250 μm CMPs. The images in exhibit A show >99% cell viability for fetal or adult, injectable CMP-MSCs/alg gel, CMP-MSCs/ml and faCMP-MSCs/bl cell sheets. The images in exhibit B show significant expansion of spindle-shape MSCs in CMPs. No significant difference in cell morphology or shape was observed between adult or fetal CMP-MSCs/alg hydrogels and CMP-MSCs cell sheets.

**Figure 10.**
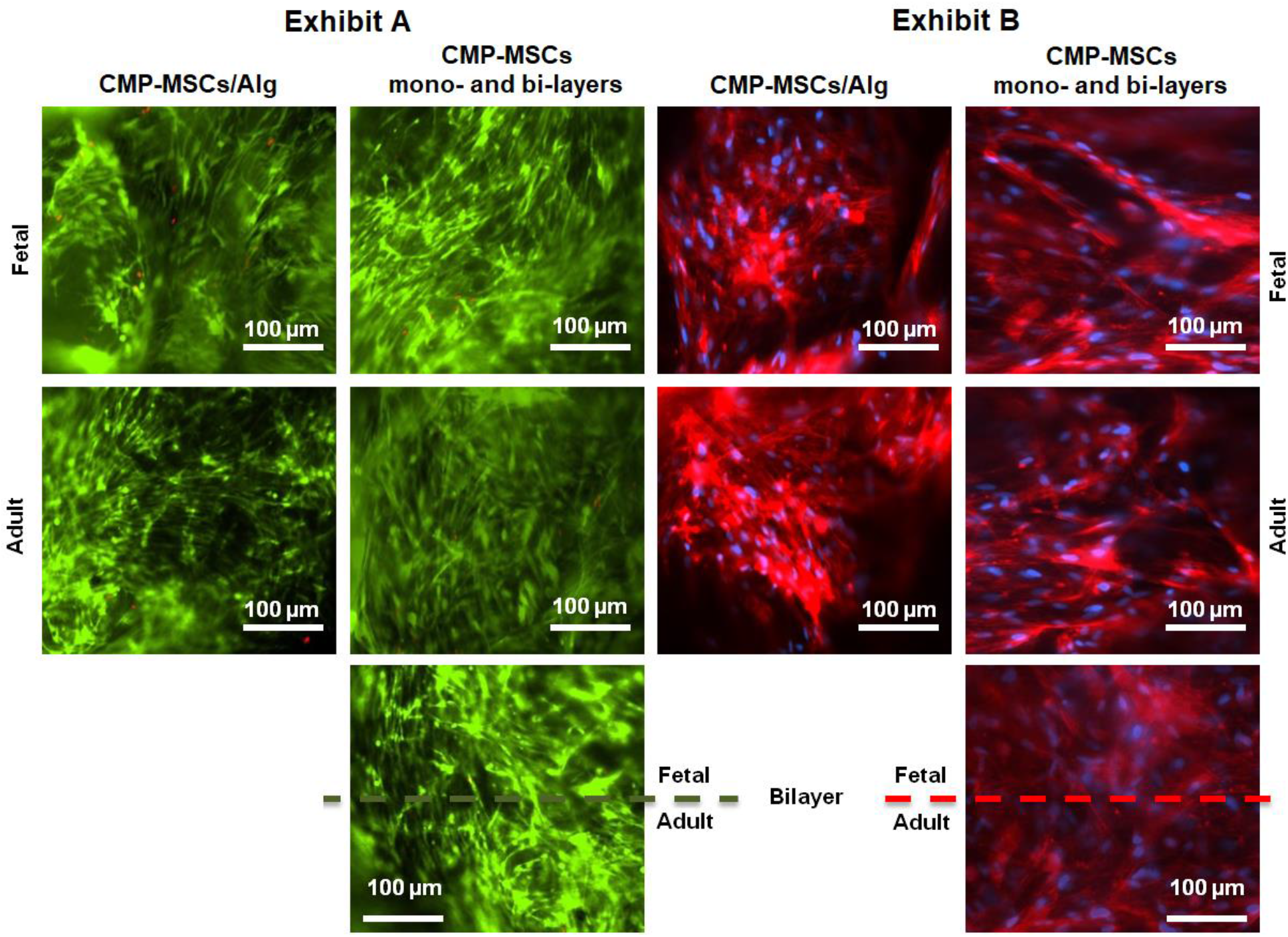
(Exhibit A) calcein AM (green) and EthD (red) fluorescent images of live and dead MSCs, respectively, in CMP-MSCs/alg hydrogels (left column) and CMP-MSCs cell sheets (right column) after 21 days incubation in chondrogenic medium; (Exhibit B) phalloidin and DAPI stained images showing cytoskeletal and nuclear compartments of MSCs, respectively, in CMP-MSCs/alg hydrogels (left column) and CMP-MSCs cell sheets (right column) after 21 days incubation in chondrogenic medium. The first and second columns in the exhibits are for fCMP-MSCs and aCMP-MSCs, respectively, and the third row is for bilayer faCMP-MSCs cell sheets. The scale bar in the images is 100 μm.

Figure 11 shows cell morphology and GAG accumulation in histological sections of adult and fetal CMP-MSCs/alg, CMP-MSCs/ml, and faCMP-MSCs/bl groups stained with H&E and Alcian blue, respectively. The fetal or adult CMPs in alginate hydrogel without MSCs were used as the control group.

**Figure 11.**
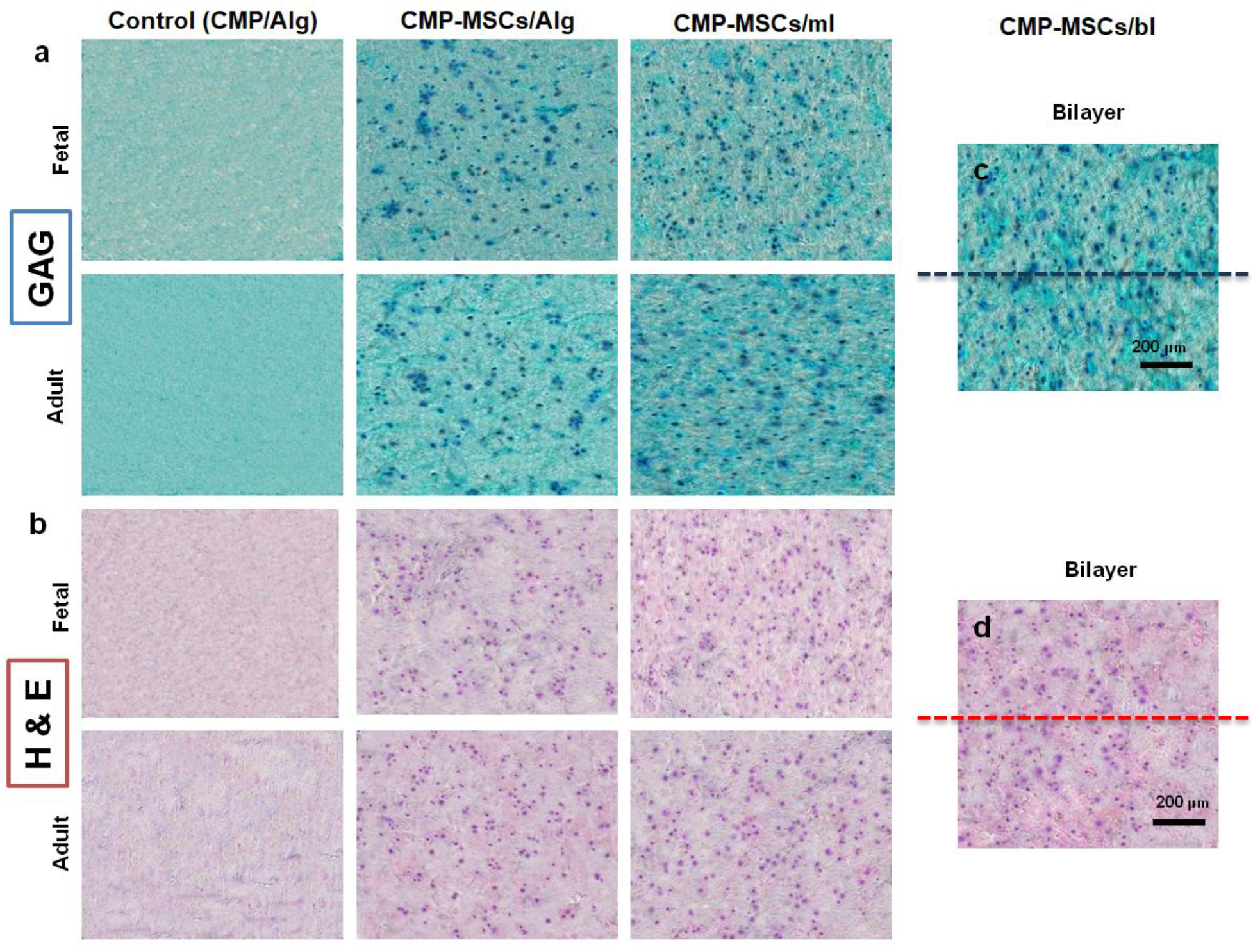
Alcian blue (a) and H&E (b) stained histological sections of control group (first column), CMP-MSCs/alg hydrogel (second column), CMP-MSCs/ml (third column) and CMP-MSCs/bl cell sheets (fourth column) after four weeks of incubation in chondrogenic medium; the first and third rows are for fetal CMPs whereas the second and fourth rows are for adult CMPs; the control group for injectable constructs was MSCs (without CMP) encapsulated in alginate and cultured in chondrogenic medium; the control group for mono- or bi-layer cell sheets was the MSC pellet cultured in chondrogenic medium. The scale bar in the images is 200 μm.

The average size of CMPs for all groups in Figure 11 was 190 μm and the CMP amount in CMP-MSCs/alg groups was 50%. There was no significant difference in cell morphology between CMP-MSCs/alg and CMP-MSCs cell sheets or between fetal and adult groups. The cell content and GAG intensity of CMP-MSCs cell sheets were slightly higher than CMP-MSCs/alg. The CMP-MSCs cell sheets had a more uniform cell and GAG distribution as compared to CMP-MSCs/alg hydrogels.

## 4. Conclusion

This work describes a novel approach to produce mono- or multi-layer cell sheets from fetal or adult articular matrix for articular cartilage tissue regeneration. Fetal or adult bovine articular cartilage was minced, decellularized, and freeze-dried. The freeze-dried ECM was grinded and sieved to produce microparticles (CMPs) with average sizes in the range of 90-250 μm. Human MSCs expanded on fetal or adult CMPs in basal medium maintained the expression of mesenchymal markers. Next, two approaches were used to generate injectable or preformed cell carriers for articular cartilage regeneration. In one approach, CMP-MSCs were suspended in alginate gel, crosslinked with calcium chloride after injection in a mold, and incubated in chondrogenic medium to generate an injectable matrix for articular cartilage regeneration. In another approach, CMP-MSCs were suspended in a culture medium, allowed to gravitationally settle on the culture plate and fuse by incubation in chondrogenic medium to generate a preformed cell sheet. Multilayer cell sheets were generated by sequential settling and fusion of zone-specific CMP-MSCs to simulate the stratified structure of articular cartilage. Fetal or adult CMP-MSCs in alginate hydrogels showed higher expression of chondrogenic markers after four weeks and compressive modulus after eight weeks of incubation in chondrogenic medium compared to MSCs directly encapsulated in alginate gel; CMP-MSCs cell sheets showed higher expression of chondrogenic markers after four weeks and compressive modulus after eight weeks of incubation in chondrogenic medium compared to MSCs in a pellet culture. The higher quality of the generated tissues for CMP-MSCs groups was attributed to superior cell-cell and cell-matrix interactions compared to MSCs encapsulated in alginate or MSC pellets. The cell sheet approach could potentially be used to produce multilayer constructs mimicking composition and cellularity of distinct zones of articular cartilage.

## Acknowledgements

This work was supported by research grants to E. Jabbari from the United States National Science Foundation under Award Numbers CBET1403545 and IIP150024 and the National Institute of Arthritis and Musculoskeletal and Skin Diseases of the National Institutes of Health under Award Number AR063745. The content is solely the responsibility of the authors and does not necessarily represent the official views of the National Institutes of Health. SK was supported by a scholarship from the Ministry of Higher Education and Scientific Research/Al-Nahrain University, Baghdad, Iraq.

